# A novel emotional and cognitive approach to welfare phenotyping in rainbow trout exposed to poor water quality

**DOI:** 10.1101/297747

**Authors:** V Colson, A Mure, C Valotaire, JM Le Calvez, L Goardon, L Labbé, I Leguen, P Prunet

## Abstract

Recent scientific evidence for fish sentience has stressed the need for novel sentience-based detection tools of fish welfare impairment in commercial farms. In order to mimic a well-characterised stress situation, rainbow trout (*Oncorhynchus mykiss*) were exposed to poor water quality (hypoxia combined with high ammonia level) for three weeks (stressed group, S) and compared to a non-stressed control group (NS). After a return to water quality standard, emotional responses were assessed in fish subjected to two potentially threatening situations: (i) social isolation in a novel environment and (ii) human presence. In addition, we used an appetitive-conditioning paradigm to determine whether previous chronic deterioration of water quality disturbs cognitive abilities in fish. Spontaneous behaviour in the tanks was also recorded during the environmental challenge as a reference for fish activity. We observed that in S fish, plasma cortisol levels were increased before and after social isolation in a novel environment compared to the plasma cortisol levels in the NS group, despite the absence of a behavioural difference between the two groups. Under deteriorated water quality, fish locomotor activity was globally reduced and this reduction was correlated to increased shoaling behaviour. Farmers can use these first behavioural modifications as a sentinel detector for fish welfare impairment. More importantly, we demonstrated that reactivity to a human presence in a home-environment and food-anticipatory behaviour were both inhibited in the S group. We consider that these two sentience-based tests are highly relevant for fish welfare assessment at the group level and are easy to use in the aquaculture industry.

## 1. Introduction

The European aquaculture industry has to cope with growing public concerns over animal welfare that lead to new regulations. With the recent scientific evidence for fish sentience (Braithwaite and Huntingford, 2004; Sneddon, 2006; Braithwaite and Boulcott, 2007; Yue et al., 2008; Kittilsen, 2013) and the recommendations concerning the welfare of farmed fish issued by the Council of Europe in 2005, there is a need for innovative sentience-based approaches to provide new standardised and practical tools that would allow farmers to easily detect any fish welfare impairment. Fish welfare studies often fail to consider the mental state of animals, although this corresponds to the fifth freedom ensuring animal welfare (FAWC, 2009). This argument implies that environmental conditions and treatments must avoid mental suffering, such as fear and distress, in animals. While extensively used in many terrestrial animals, welfare markers based on emotional states are still very scarce in farmed fish, despite the fact that fish have sufficient neural capacities to potentially support the experience of emotion and complex cognitive abilities (see Braithwaite et al. (2013) for a review). In agreement with Mendl (2001), we consider the measurement of emotional reactions to be crucial in assessing animal welfare state. Indeed, emotional responses are more likely determined by the way the animal appraises its environment than by the environment itself (Mason, 1971; Boissy et al., 2007; Boissy and Erhard, 2014), and this appraisal can vary according to the previous or present living conditions. For instance, emotional behavioural responses are known to be modified after chronic stress, induced by unpredictable aversive events in mammals (Boissy et al., 2001; Destrez et al., 2013), birds (Calandreau et al., 2011) and fish (Piato et al., 2011). Poor environmental conditions (e.g. barren cages) are also associated with a stronger response to novelty and increased fearfulness in laying hens (Whay et al., 2007; Nicol et al., 2009). Measuring the increase in fearfulness in acute challenging situations could therefore be used to assess the chronic stress state in farmed fish. In animals, emotional reactions can be inferred from acute behavioural and physiological adjustments. Behavioural adjustments in response to alarming situations reflect fearfulness, and can be observed either through proactive reactions (e.g. flight, aggression) or reactive (i.e. passive) reactions (e.g. freezing, hiding). These different behavioural patterns have been described in rainbow trout (*Oncorhynchus mykiss*) selected for high- and low-stress responsiveness (Overli et al., 2002). Fish fearfulness can also be evaluated through the monitoring of plasma cortisol levels, a classical stress response marker (Pottinger and Mosuwe, 1994; Mommsen et al., 1999; Mormede et al., 2007). In the present study, we tested an emotional approach for fish welfare phenotyping, based on these behavioural and cortisol parameters, in response to a challenge of social isolation in a novel environment.

Stress is also known to have a cognitive impact in many animal species. Exposure to acute or chronic stressors usually leads to impaired learning and memory in mammals (Diamond et al., 2006; Wood et al., 2008; Green and McCormick, 2013). Interestingly, spatial learning abilities can be improved in quails, after one week of stress (Calandreau et al., 2011). Farmed fish can easily learn to stay or move to a specific location in the tank in order to get food (Bratland et al., 2010; Martins et al., 2012). This food-anticipatory activity can be a sensitive measure for assessing the effect of chronic stress on cognition. Moreover, the anticipatory state reflects the need and the inclination of an animal for pleasant events (e.g. meals), and represents, by itself, a good approach for observing welfare from the perspective of the animal (Spruijt et al., 2001). In this study, we measured fish learning rate in a classical conditioning paradigm at the group level to easily evaluate the impact of chronic stress not only on cognition but also on the general inclination of the fish for pleasure (e.g. positive emotion), by anticipating a food reward.

The purpose of this study was to develop several innovative and complementary emotional and cognitive approaches for welfare phenotyping that may, in the future, be used under aquaculture conditions. To this aim, deteriorated water quality (hypoxia combined with high ammonia levels) induced by reducing water renewal in tanks was used as a chronic stressor in a case study, reflecting common problems encountered in aquaculture. Under commercial farming conditions, deterioration of water quality can have dramatic negative effects on fish food intake, food conversion efficiency, growth (Person-Le Ruyet et al., 2008; Santos et al., 2010), health (Cohen et al., 2012), body injuries (d’Orbcastel et al., 2009), and physiology and endocrinology (Caldwell and Hinshaw, 1994; Vianen et al., 2001; McNeill and Perry, 2006; O’Connor et al., 2011). Several effects on fish behaviour have also been observed in response to hypoxia. Fish subjected to hypoxia display increased surface aquatic respiration (Abdallah et al., 2015), decreased locomotor activity (Herbert and Steffensen, 2005; Xu et al., 2006; Espmark and Baeverfjord, 2009), and altered anti-predator behaviour, with disruption of the school unit (Domenici et al., 2007). Rainbow trout is a major aquaculture species and chronic deterioration in water quality has been demonstrated as being critical for welfare, as evaluated by physiological, health and growth classical welfare indicators (Caldwell and Hinshaw, 1994; vanRaaij et al., 1996; Vianen et al., 2001; Person-Le Ruyet et al., 2008).

In the present study, behavioural responses reflecting mental states were assessed in rainbow trout chronically exposed to deteriorated water quality and compared to a control group. For this aim, fish were subjected to 2 potentially threatening situations: social isolation in a novel environment and human presence. In addition, we used a food-anticipatory paradigm to determine whether chronic exposure to poor water quality prevents the formation of a reward-location association in rainbow trout. Spontaneous behaviour was also recorded in the tanks as a reference for fish activity.

## 2. Material and methods

### 2.1 Ethics statement

Experiments were conducted within the INRA-PEIMA (Sizun, France) facilities with authorization for animal experimentation (C29-447-02). All experimental procedures used in this study complied with the rules framed by the “National Council for Animal Experimentation” of the French Ministry of Higher Education and Research and the Ministry of Food, Agriculture, and Forest. The poor water conditions used in this study were at the tolerance limits for this species. Such conditions were needed to create a relevant chronic stressor. The minimum dissolved oxygen concentration registered during the 3-week hypoxic period was still above the minimum critical oxygen level considered to be lethal in rainbow trout (Pennell and Barton, 1996), and would be experienced in natural pools, particularly during warmer summer months (Matthews and Berg, 1997). Therefore, although these factors might affect metabolic processes, the fish are known to acclimate to and tolerate such conditions. The present project was approved by the local “Ethic, Animal Care and Use Committee” provided by the French legislation under the official license N°74. The project’s agreement number is: 02533.01.

### 2.2. Fish and chronic stress protocol

This study was carried out using immature triploid rainbow trout (Autumn-spawning ©INRA strain), since most rainbow trout production in Europe is based on sterile triploid fish. Note that triploid fish are less tolerant than diploids to chronic hypoxia (Hansen et al., 2015). However, recent evidence indicates that there is no major difference in the cortisol stress response between diploid and triploid fish (Fraser et al., 2012). Breeding and experiments were performed at PEIMA (INRA fish farming, Sizun, France), in an open flow through system supplied with river water. When fish were 9 months old (i.e. 272 days post-fertilisation), they were distributed between 12 identical grey resin tanks (1 m x 1 m x 0.2 m depth) and acclimatized to a 50 kg/m^3^ density (87-89 fish per tank supplied with 200 l of water) for 2 weeks. Flow rate was approximately 700 l of water per hour. Following this acclimation period, the flow rate was monitored for all tanks over a period of 21 days (D0 to D21), and adjusted to produce the 2 experimental groups (n = 6 tanks / group) with differing levels of dissolved oxygen: the stressed group, S (poor water conditions, O_2_: min-max = 3.1-4.9 mg/l), and the non-stressed control group, NS (standard conditions, O_2_: min-max = 8.0-11.0 mg/l). Global mean weight was measured for each tank in S and NS groups at the start of the acclimation period to ensure no difference existed initially between groups (mean weight ± SEM: 113.6 ± 0.3 g and 114.4 ± 1.3 g, respectively). These initial weights were not significantly different (Wilcoxon non-paired test: W = 26, *P* = 0.23).

In the stressed group, at D0, D1, D2, D3, D9, D11, D12, D13, D14, D15, D17, D18 and D21 (once each day between 13:00 and 17:00), the dissolved oxygen levels in the effluent water of each tank were recorded using a portable probe connected to an Optical Density Reader (Hach-Lange, France). In the control group, oxygen monitoring was carried out at D0, D2, D3, D11, D12, D13, D14 and D21. The oxygen levels over the 3-week period are reported in Fig. 1. To attain the required experimental oxygen level in each treatment, the experimenter activated the tap adjusting the flow rate in each tank (stressed group: flow ranging between 144 and 180 l/h; control group: flow ranging between 612 and 720 l/h). In each tank, the water inlet was positioned at the opposite side of the feeder and a submerged pipe distributed the inflowing water towards the tank wall, causing a circular flow pattern. The outlet was positioned in the centre of the floor of the tank. Ammonia levels were measured on D7 and D14 in 2 control and 2 stressed tanks (the same 4 tanks on each day) (stressed group: ammonium nitrogen (NH4-N) = 0.71 ± 0.01 mg/l and 0.56 ± 0.01 mg/l, respectively; control group: NH4-N = 0.35 ± 0.03 mg/l and 0.37 ± 0.03 mg/l, respectively). Water pH was 6.5. At this pH, ammonia was in the ionized form, which is the less toxic form of ammonia. The temperature ranged between 10°C and 12°C during the experimental period. Ammonium nitrogen was measured with an ammonia test kit (Hach-Lange, France). Throughout the 3-week challenge period, we recorded a total of 11 dead fish from the 529 fish initially reared in the stressed condition, i.e. 2% mortality (*vs* 0% in the control condition).

**Figure 1.**
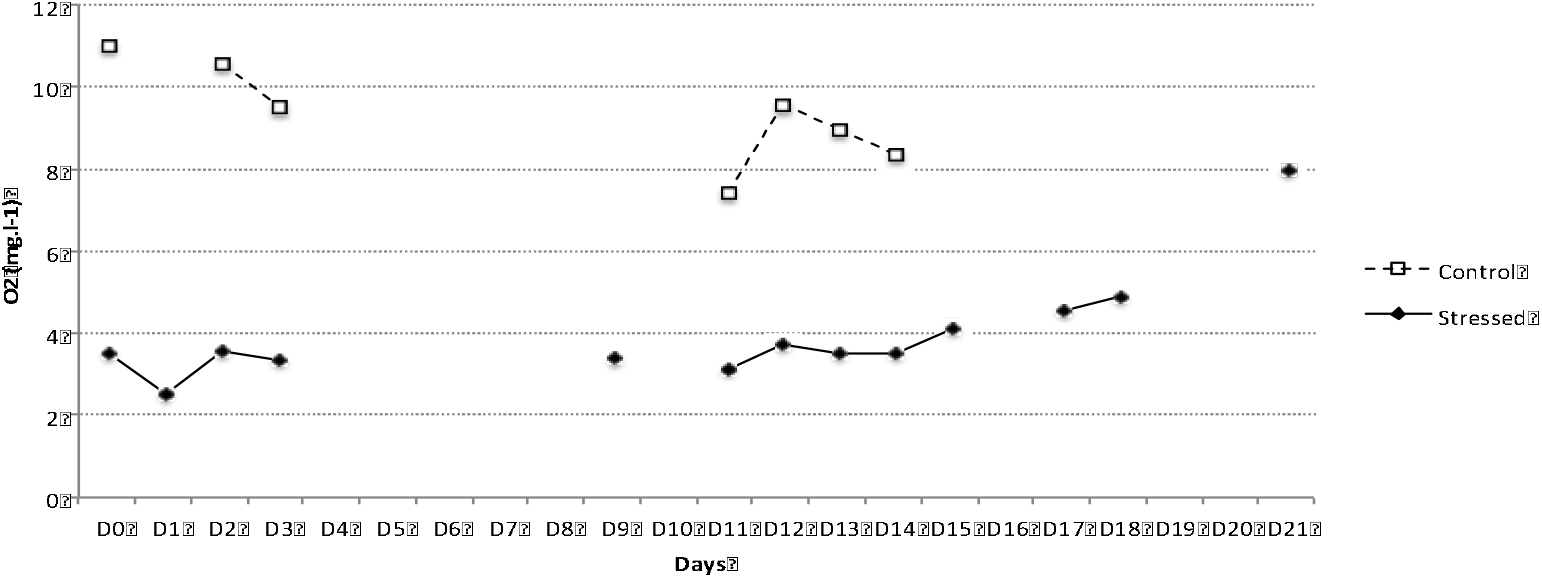
Mean (± SEM) oxygen levels measured in the Control group (n = 6 tanks) and in the Stressed group (n = 6) between D0 and D21 in the afternoon (between 13:00 and 17:00).

Fish were fed 3 times per day *ad libitum* with a commercial diet (43% proteins and 24% lipids), supplied by Arvotec® automatic feeders. Fish were starved for 24 h before individual behavioural tests.

A 30 cm-vertical net prevented the fish from escaping during acclimation. Tanks were equipped with digital cameras positioned directly above the tanks (one camera for two tanks). Luminosity was controlled and the room was automatically illuminated from 08:00 to 20:00.

### 2.3. Behavioural parameters

#### 2.3.1. Focal observations

After the 2-week acclimation period, behavioural parameters were video-recorded for 10 minutes in both NS and S groups (6 non-stressed and 6 stressed tanks) at Day-1 (the day before the start of chronic stress), Day+1, D4, D11, D16 and D18, at 1600h (before the last meal). Using the continuous sampling method (when all of the activities that occur in a group of individuals are recorded for a specified time period), the following behaviours were recorded for each tank: stereotypies (repetitive swimming against the edge of the tank), acceleration bursts, and jumps. In addition, scan samplings (1/minute) (when the behaviour of all the individuals in a group are recorded at predetermined time intervals) were performed to measure the time spent inactive (as described in Colson et al. (2015b)) and the time spent grouped as a shoal. The results are presented as percent of the observed time (over 10 scans) the fish were ‘inactive’ or ‘grouped’. Behaviours are thoroughly described in Table 1.

**Table 1.** Behavioural parameters used for focal observations (10 minutes/day)

| Behaviour | Observation method | Definition |
| --- | --- | --- |
| stereotypies | focal sampling (occurrence) | Repetitive rubbing of the mouth against the wall of the tank with head emerged |
| jump | focal sampling (occurrence) | Movement towards the water surface with sudden emergence of at least half of the body out of the water |
| acceleration bursts (eg fast-start) | focal sampling (occurrence) | Sudden movement away from the initial position with propulsive tail strokes without agonistic behavior |
| inactive | enlarged scan sampling (10 sec/min) | No displacement of the fish among at least two thirds of the group in the tank. Results are expressed as % of scans (over 10 scans) this phenomenon was observed |
| grouped | scan sampling (1 scan/min) | Aggregation of fish as a shoal that occupied less than half of the tank. Results are expressed as % of scans (over 10 scans) this phenomenon was observed |

#### 2.3.2. Open-field test: behaviour and cortisol

Twenty-four hours following the return to standard quality water (on D22 and D23), 12 fish per experimental condition were subjected to an open-field test, as previously described in rainbow trout (Schjolden et al., 2005; Colson et al., 2015a). In each condition, we used a total of 4 fish per tank, with 3 tanks dedicated to this test, and the 3 remaining tanks dedicated to the basal cortisol sampling, which occurred prior to the test (see below). This experiment consisted of netting 4 fish simultaneously and immediately transferring them into 4 novel tanks (1 × 1 × 0.80 m), positioned close to the initial rearing tanks, and having the same water quality parameters. This operation was repeated to obtain 12 fish per experimental condition. The experimental condition order was chosen at random. Immediate behavioural responses of isolated individuals were video-recorded for 20 minutes (with a sampling rate of 5 pictures per second) and analysed using EthovisionXT® software (Noldus, Wageningen, The Netherlands). The tank containing the experimental fish was divided into two areas, a central zone (0.6 × 0.6 m) and a 20-cm border zone. For each individual, mean and maximum swimming speed (body length/s), percentage of time spent in thigmotaxis (e.g. in the border zone) and time spent moving (starting when velocity was higher than 0.1 body length/s) were recorded. Due to poor video quality, 2 individuals from the S group were removed from the behavioural analyses.

Before the open-field test, blood was collected from 12 fish per condition, for basal cortisol level measurements. Within each experimental condition, 4 fish per tank were collected from 3 dedicated tanks. The body weight (W) and length (L) were measured post-mortem following each blood sample. For each fish, the condition-factor was calculated as followed: K-factor = 100 (W/L^3^)

Blood samples, used for final cortisol levels, were collected from each tested trout 30 minutes after the beginning of the test. The fish were netted and transferred to buckets containing tank water with 2-phenoxyethanol (0.1% total volume). Blood was sampled from the caudal sinuses into heparinized syringes and samples were stored on ice. After sampling, blood cells and plasma were separated by centrifugation (8 minutes at 1500 rcf). Plasma was collected and frozen at −20°C until analysis. Before the cortisol assay, steroids were extracted twice from 50 µl plasma with 2 ml ethylacetate/cyclohexane. After solvent evaporation, steroids were suspended in 250 µl assay buffer. Cortisol was assayed in duplicate by a 3H cortisol radioimmunoassay as previously described (Auperin et al., 1997; Colson et al., 2015b).

#### 2.3.3. Human avoidance test

On Day 24, the fish group response to human approach was assessed for all tanks (6 control and 6 stressed tanks) using the same experimenter. 2.5 min after the start of video recording, the experimenter, wearing a red coat, approached to within 20 cm of each tank in turn, where he remained for 5 minutes until the end of the video recording.

Videos were manually analysed and fish flight response to human approach was assessed by scan sampling every minute. At each scan, we noted whether the fish were (i) horizontally distributed covering the whole tank, (ii) half grouped, or (iii) all grouped as a shoal at the opposite side (scores: 1 to 3, respectively).

#### 2.3.4. Appetitive conditioning

On Days 29, 30 and 31 (i.e. 2 weeks following the end of the chronic environmental stressor), food-anticipatory behaviour was assessed within both experimental conditions using a group conditioning procedure. As for the human avoidance test, all tanks were used for this test (6 per experimental condition), with each tank containing the remaining fish (83-85 fish / tank). The acquisition phase consisted in exposing fish to a neutral stimulus (Conditioned Stimulus, CS: water-flow off and on) 6 times a day for 3 days and to feed them 60 seconds later (Unconditioned Stimulus, US: Food-reward). Every stimuli association was video recorded for 120 seconds, with the CS and US occurring at 60 and 120 seconds respectively. The pump took 20 seconds to stop the flow and 20 more seconds were needed to turn the pump on again. The water flow was completely recovered after 20 more seconds. Thus, the CS took a total of 60 seconds, after which the US immediately occurred. Note that the water-flow offset represented a neutral signal very distinct from the permanent low current used to deteriorate water quality during the 3-week of environmental stressor. At each trial (from 1 to 18), the conditioned response was evaluated by scan samplings performed at 0, 65, 75, 105 and 120 seconds. At each scan, a learning index was given ranging from 1 to 4 (1: fish were horizontally distributed throughout the whole tank; 2: less than 1/3 of the fish were grouped in the feeding area; 3: half of the fish were grouped in the feeding area; 4: more than 2/3 of the fish were grouped in the feeding area with scramble competitions). This index is more sensitive than that of the human approach test since differences between groups are subtler and the intervals between scans are shorter in a conditioning test.

For the behavioural parameters obtained during the open-field test and for body weight and length measurements, Student’s t-tests were used to compare the two experimental groups, since data distribution reached normality (evaluated by a Shapiro–Wilk normality test). For cortisol levels, a two-way ANOVA was performed to determine the effects of chronic stress (2 levels), acute stress (i.e. before and after the open-field test, 2 levels) and the interaction between these two fixed factors. The significance of the differences between means was tested using post-hoc tests (Tukey’s HSD test).

For focal observations, two-way ANOVAs were performed to determine the effects of chronic stress (2 levels), day (paired data, 5-6 levels) and their interaction (n = 6). The significance of the differences between means was tested using post-hoc tests (Tukey’s HSD). In order to reach a normal distribution in behavioural data (Shapiro–Wilk normality test), quantitative data were square-root transformed and the proportions (0-1) of time spent immobile or grouped (focal observations), in thigmotaxis or moving (open-field test) were arc-sine-square-root transformed.

For appetitive conditioning and human approach data, scores did not reach normal distribution, even after transformation. Therefore, we performed the non-parametric Mann-Whitney test (R, Mann-Whitney-Wilcoxon test) at each time point between the two independent groups. Between trials comparisons were performed within each treatment using Wilcoxon tests since trials were dependant data.

To examine the relationship between time spent immobile and time spent grouped, we used Pearson’s correlations on asin transformed data.

Differences were found to be significant when P < 0.05. The analyses were executed using the free software R 2.14.0 (http://cran.r-project.org/).

## 3. Results

### 3.1. Body weight and length

At the end of the experiment, body weight was significantly higher in control fish (mean weight ± SEM = 188.25 ± 8.22 g) than in stressed fish (153.9 ± 7.65 g; t = 5.45, df = 36.82, P < 0.001; Table 2). Additionally, control fish were significantly longer than stressed fish (mean lengths ± SEM: 24.2 ± 0.28 cm *vs* 23.34 ± 0.27 cm; t = 3.94, df = 38.21, P < 0.001). The condition-factor was also significantly different between control and stressed fish (mean K-factor ± SEM: 1.32 ± 0.03 *vs* 1.20 ± 0.02; t = 5.13, df = 39.93, P < 0.001).

**Table 2.** Mean (± SEM) weight (W), length (L), condition factor (K-factor = 100 x W/L^3^) and plasma cortisol levels measured on control and stressed fish 30 minutes after the start of the open-field test. Behavioural parameters (mean ± SEM) were observed during the 20-minute test. Control and stressed groups were compared using Student t-tests.

|  | Control | Stressed | n (Control) | n (Stressed) | P-value |
| --- | --- | --- | --- | --- | --- |
| Weight (g) | 188.25 ± 8.22 | 153.9 ± 7.65 | 25 | 24 | < 0.001 |
| Length (cm) | 24.2 ± 0.28 | 23.34 ± 0.27 | 25 | 24 | < 0.001 |
| K-factor | 1.32 ± 0.03 | 1.20 ± 0.02 | 25 | 24 | < 0.001 |
| Plasma cortisol (ng/ml) | 35.52 ± 4.09 | 57.73 ± 4.05 | 12 | 12 | < 0.05 |
| Thigmotaxis (% time) | 78.54 ± 7.08 | 82.42 ± 5.13 | 12 | 10 | ns |
| Movement (% time) | 38.87 ± 10.04 | 34.16 ± 6.99 | 12 | 10 | ns |
| Mean swimming speed (BL/s) | 8.43 ± 2.20 | 7.34 ± 2.07 | 12 | 10 | ns |
| Maximum swimming speed (BL/s) | 3.40 ± 1.85 | 4.15 ± 2.41 | 12 | 10 | ns |

### 3.2. Focal observations

Stressed groups had lower levels of acceleration bursts (2-way ANOVAs, F_1,70_ = 21.36, P < 0.001; Fig. 2a), stereotypies (F_1,70_ = 61.45, P < 0.001; Fig. 2b) and jumps (F_1,76_ = 9.53, P < 0.01; Fig. 2c). A day-effect was observed over time with decreasing accelerations (F_6,70_ = 3.71, P < 0.01), stereotypies (F_6,70_ = 14.9, P < 0.001) and jumps (F_6,76_ = 3.34, P < 0.01). The interaction between treatments and days was significant for the first 2 items, respectively (F_6,70_ = 3.36, P < 0.01; F_6,70_ = 3.97, P < 0.01), but not for the number of jumps (P = 0.2).

**Figure 2.**
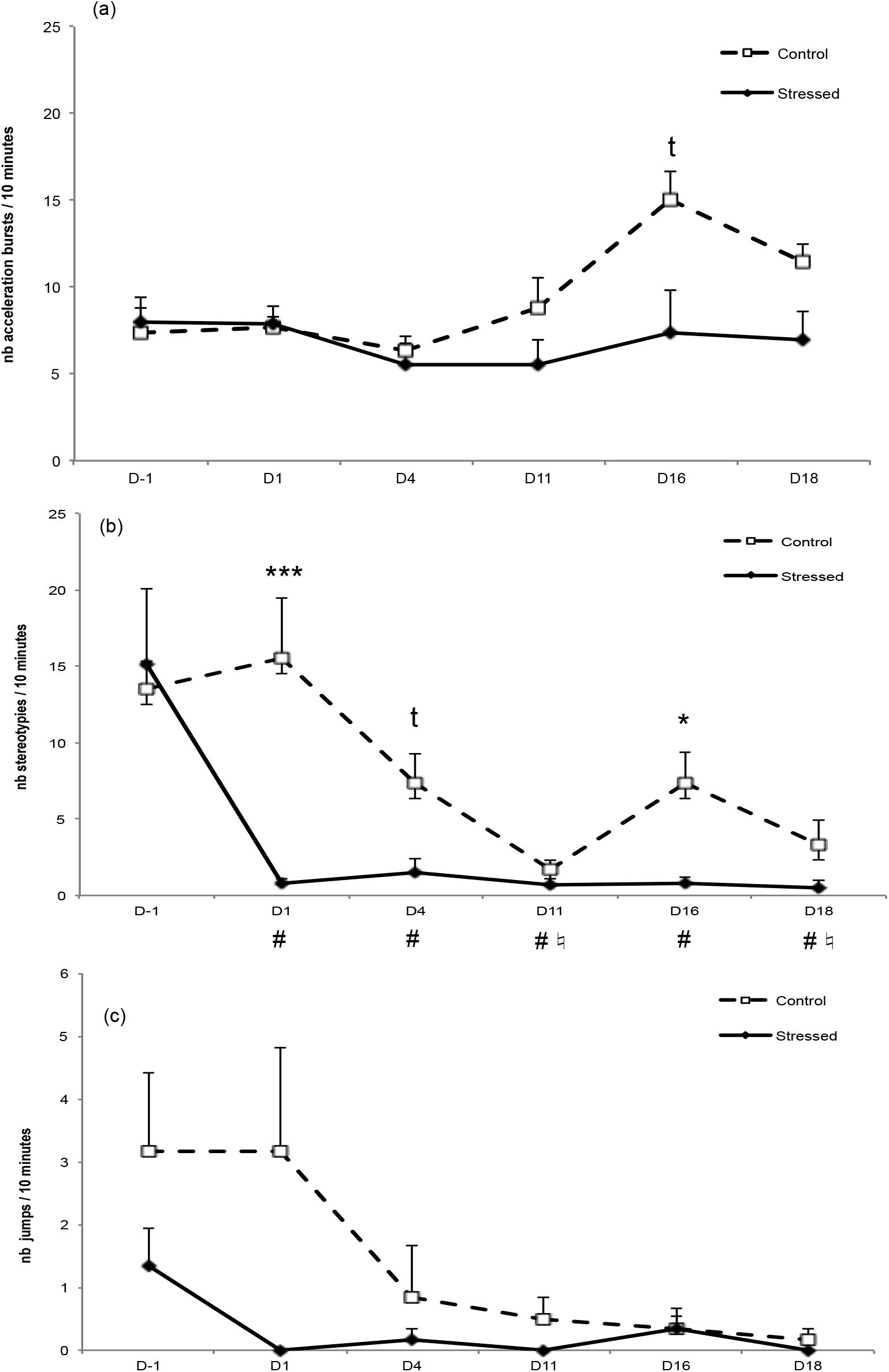
Mean (± SEM) occurrence of acceleration bursts (a), stereotypies (b) and jumps (c) observed from Day-1 to Day18 during a 10-minute focal sampling in control and stressed rainbow trout (n = 6). Asterisks denote a significant difference between control and stressed fish (*** P < 0.001, * P < 0.05, t 0.05 < P < 0.1). LJ means a significant difference between DX and D-1 in the stressed group (P < 0.01) and ♮ means a significant difference between DX and D-1 in the control group (P < 0.01).

Post-hoc Tukey’s tests showed that the number of acceleration bursts tended to be higher in the control group than in the stressed group at D16 (P = 0.058). The number of stereotypies was higher in the control group at D1 (P < 0.001) and D16 (P = 0.01).

In the stressed group, a significant decrease in stereotypies was observed between D-1 and all subsequent days (P < 0.001). In the control group, a decrease in stereotypies appeared from D1, with significant differences between D1 and D11 (P < 0.001) as well as D1 and D18 (P < 0.05).

Regarding the amount of time spent inactive, significant effects of the treatment (2-way ANOVA, F_1,50_ = 56.22, P < 0.001; Fig. 3a), days (F_4,50_ = 12.55, P < 0.001) and their interaction (F_4,50_ = 10.95, P < 0.001) were observed. Post-hoc tests showed a higher amount of time spent inactive in the stressed group at D11 (P < 0.001), D16 (P < 0.001) and D18 (P < 0.01). The increase between D-1 and D11, D16, and D18 was significant (P < 0.001).

**Figure 3.**
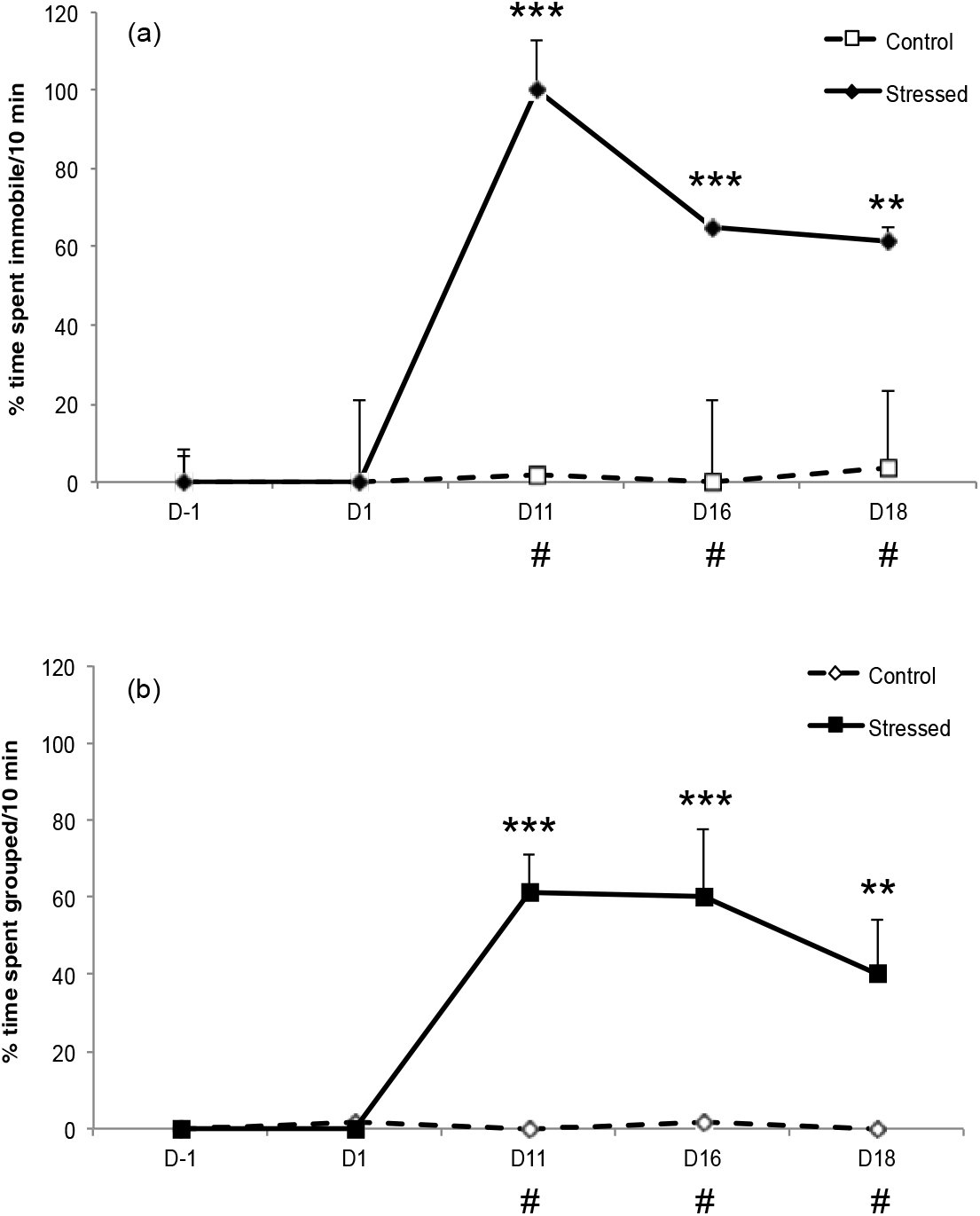
Mean (± SEM) % of time spent immobile (a), and grouped (b) observed from Day-1 to Day18 during a 10-minute focal sampling in control and stressed rainbow trout (n = 6). Asterisks denote a significant difference between control and stressed fish (*** P < 0.001, ** P < 0.01). LJ means a significant difference between DX and D1 in the stressed group (P < 0.01).

We found significant effects of the treatment (F_1,50_ = 49.77, P < 0.001), days (F_4,50_ = 9.59, P < 0.001) and their interaction (F_4,50_ = 9.79, P < 0.001) on the amount of time that was spent grouped (Fig. 3b). Stressed fish spent significantly more time grouped than control fish at D11 (P < 0.001), D16 (P < 0.001) and D18 (P < 0.01). The increase between D-1 and D11, D16, and D18 was significant (P < 0.001). A positive Pearson’s correlation was found between the time spent immobile and the time spent grouped as a shoal (t = 11.07, df = 82, correlation coefficient = 0.77, P < 0.001).

### 3.3. Open-Field test

Behavioural data

#### 3.3.1 Behavioural data

No significant difference was observed between control and stressed fish in any behavioural parameter (Table 2).

#### 3.3.2. Cortisol data

Cortisol levels were significantly increased by the open-field test (F_1,41_ = 86.50, P < 0.001; Fig. 4). Post-hoc Tukey’s HSD comparisons showed significant differences between baseline and final levels in both groups (P < 0.001). Treatment had a significant effect on cortisol levels (F_1,41_ = 18.37, P < 0.001). Before the acute stress, the cortisol levels in stressed-fish tended to be higher than the levels in control fish (Tukey’s HSD: P = 0.05), and this difference was significant after the acute stress (control: 35.52 ± 4.09 ng/ml *vs* stressed: 57.73 ± 4.05 ng/ml, P < 0.001). No interaction effect was found between treatment and acute stress (F_1,41_ = 1.42, P = 0.24).

**Figure 4.**
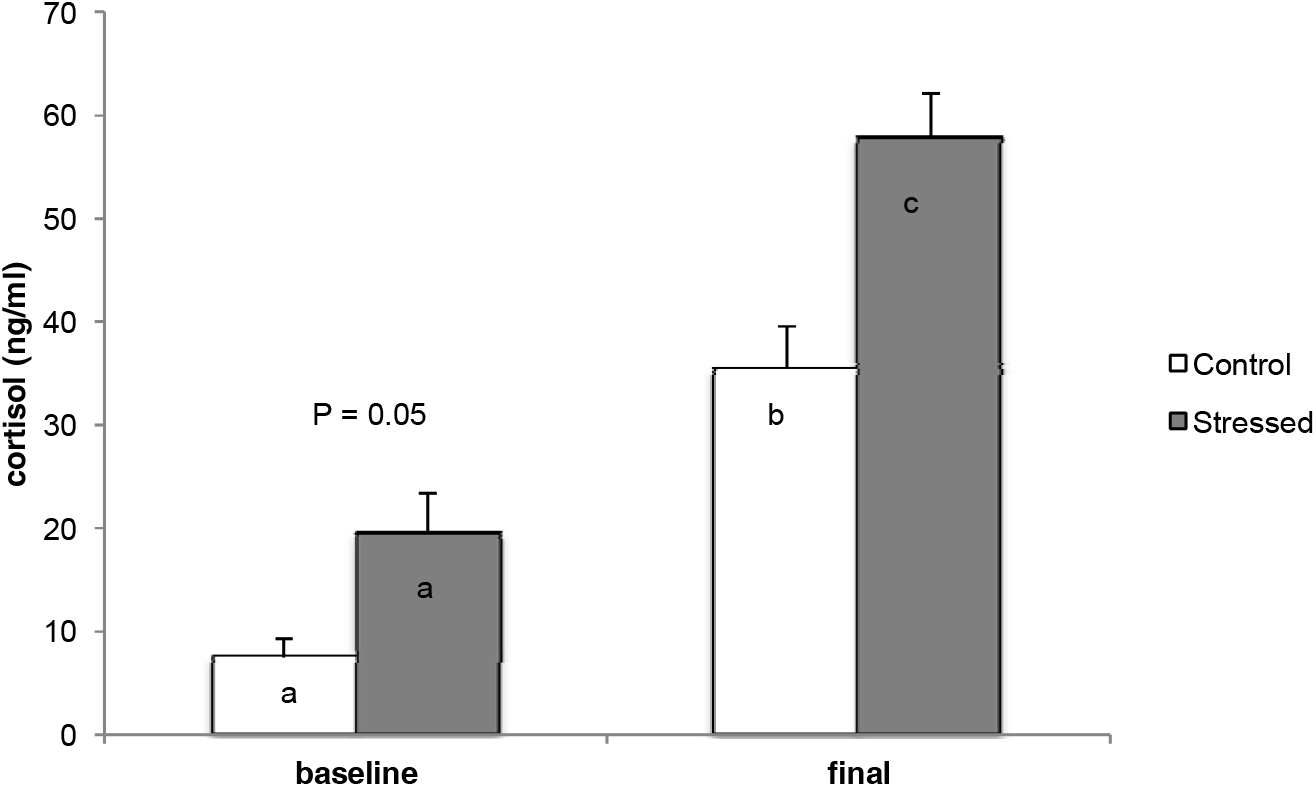
Mean (± SEM) plasma cortisol levels in control (n = 12) and stressed fish (n = 12) before and after the open-field test. The effects of the treatment, the open-field test and the interaction treatment X test were evaluated by a two-way ANOVA, followed by Tukey-HSD tests. Statistical differences between treatments are denoted by different letters (P < 0.001).

### 3.4. Human avoidance test

Fish from both groups were similarly dispersed before the human avoidance test. When the human approach occurred (minute 4), fish from the control tanks were more grouped as a shoal than fish from stressed tanks (Mann-Whitney test: W = 33, n = 6, P < 0.01, Fig. 5).

**Figure 5.**
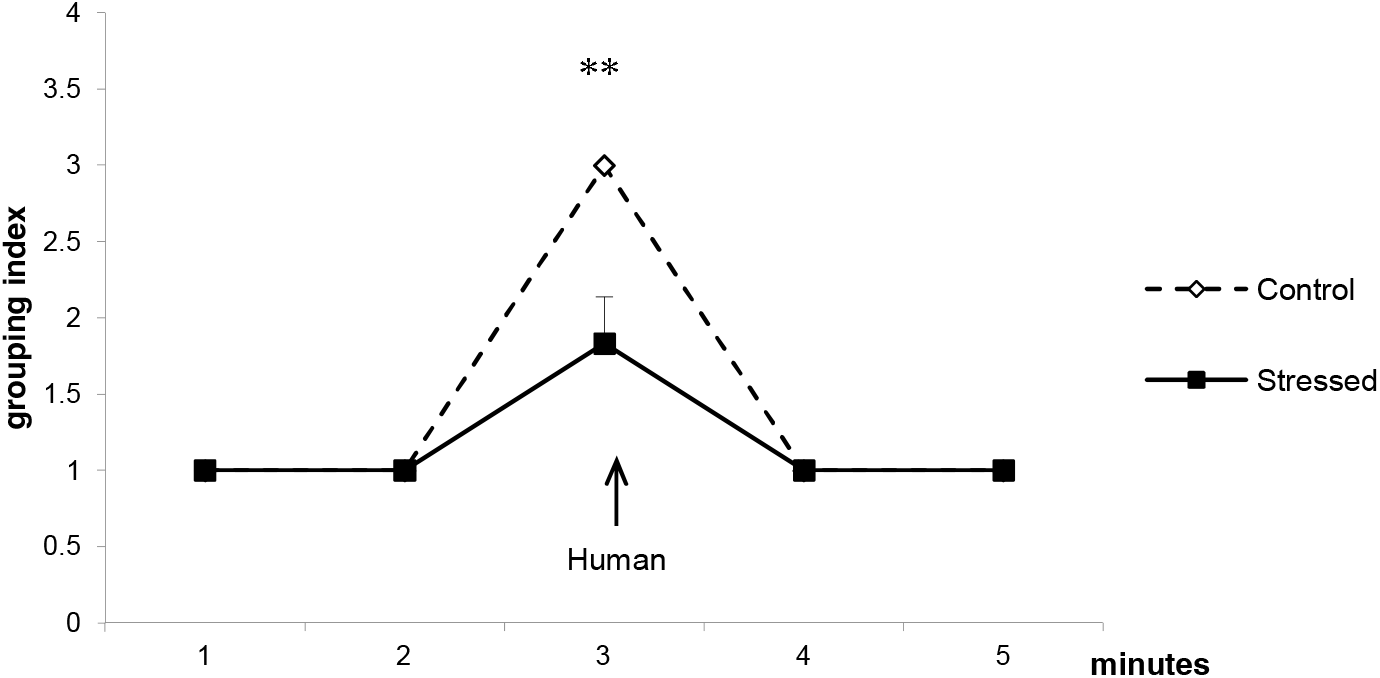
Mean (± SEM) grouping index of control and stressed groups (n = 6) during the Human-avoidance test. Mann-Whitney-Wilcoxon tests at each time point were used. ** P< 0.01.

### 3.5. Appetitive conditioning

On trial 1, the conditioning index obtained at each scan did not differ between the control and the stressed groups (Fig. 6a). On trial 18, fish from the control group had a higher conditioning index than fish from the stressed group when the CS occurred at scan 65 (Mann-Whitney test: W = 30, n = 6, P = 0.028, Fig. 6b) and at scan 75 (W = 30, n = 6, P = 0.027). The difference disappeared at scan 105 (W = 20.5, P = 0.67). When the reward (US) was given at scan 120, fish from both treatments were all grouped in the feeder area.

**Figure 6.**
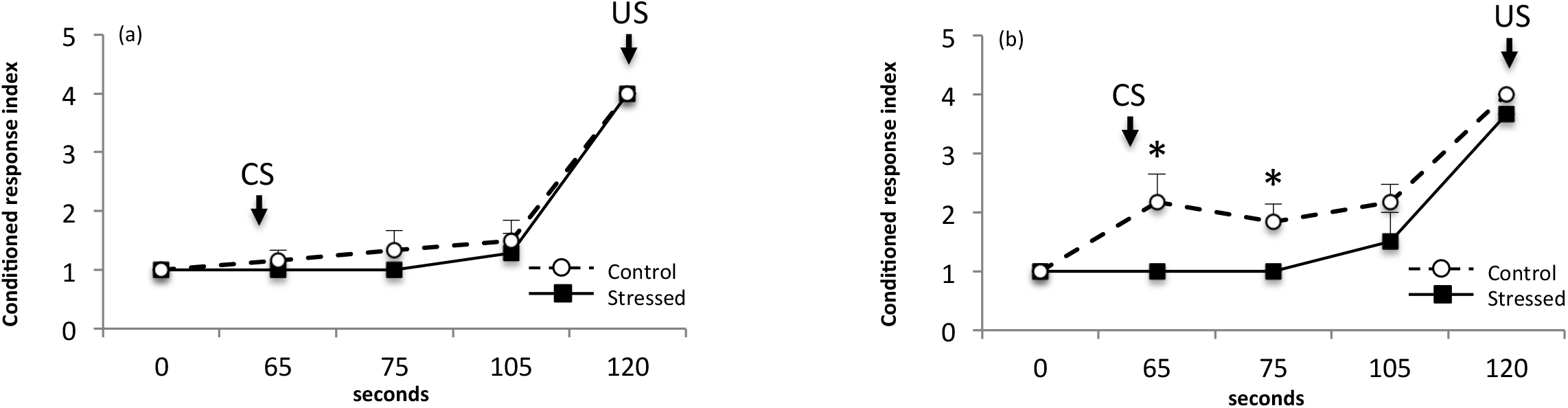
Mean (± SEM) conditioned response index of control and stressed groups (n = 6) at (a) Trial 1 and (b) Trial 18. CS (Conditioned Stimulus): Water-flow stopped at 60 seconds. US (Unconditioned Stimulus): Food-reward occurred at 120 seconds. Mann-Whitney-Wilcoxon tests at each time point were used. * P < 0.05.

At time-point 65 seconds (ie 5 seconds after the offset of the water-flow), the conditioned response index was higher in the control group than in the stressed group at trial 18 (W = 30, n = 6, P = 0.028, Fig. 7a). A tendency appeared at trial 5 and trial 6 (W = 27, P = 0.07). No significant increase of the conditioned response within any group was observed across trials at this time-point.

**Figure 7.**
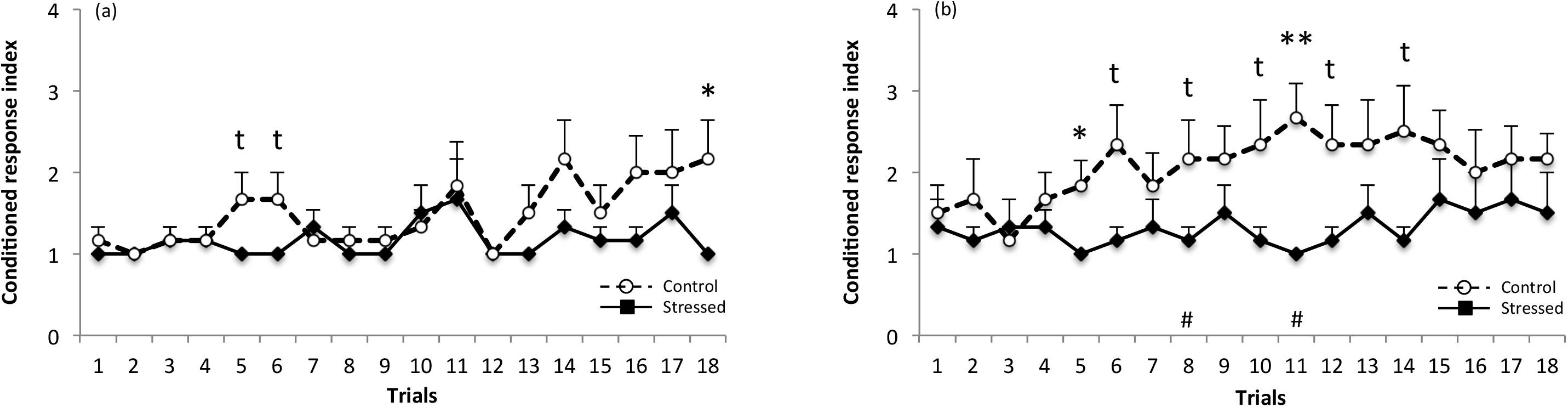
Mean (± SEM) conditioned response index of control and stressed groups (n = 6) at (a) time-point 65 seconds (i.e. 5 seconds after the offset of the water-flow), and (b) time-point 105 seconds (i.e. 5 seconds after the onset of the water-flow and 15 seconds before food-reward).

At time-point 105 seconds (ie 5 seconds after the onset of the water-flow and 15 seconds before the food-reward), the conditioned response index was higher in the control group than in the stressed group at trial 5 (W = 30, n = 6, P = 0.027, Fig. 7b) and trial 11 (W = 33, P < 0.01). Conditioned index tended to be greater in the control group at trial 6, trial 8, trial 10, trial 12 and trial 14 (0.05 < P < 0.1). Regarding the progress of the conditioned response between trial 1 and the subsequent trials, an increase tended to be significant in the control group only (trial 1 *vs* trial 8: Wilcoxon test for paired data: V = 0, P = 0.07 and trial 1 *vs* trial 11: V = 0, P = 0.09). In the stressed group, no increase was observed in the conditioned response across trials.

## 4. Discussion

A chronic exposure to poor water quality, induced by reducing water renewal, was performed in farmed fish in order to reveal novel sentience-based welfare markers that would be easy to use in husbandry systems. The deterioration in water quality included hypoxia, ammonia and possibly the increase in total suspended solid concentration. These experimental conditions were representative of poor water quality encountered in commercial rearing systems. However, the elevation of ammonium nitrogen observed in our tanks remained significantly below the recommended thresholds for rainbow trout (Thurston et al., 1981).

### Fish behaviour observed by focal observations

Oxygen is a limiting factor for metabolism, and salmonids show reduced growth at levels that are considered mild hypoxia (70% saturation at 16°C; Remen et al., 2012). Here, severe hypoxia was applied (40% saturation at 10-12°C). This explains important morphological (weight, length and K-factor) discrepancies between stressed and control groups after 3 weeks. The overall focal observations indicate that the chronic stress protocol dramatically reduced fish group activity. This is in accordance with various previous studies, which also demonstrated a decrease in locomotor activity in different fish species exposed to hypoxic conditions (goldfish (*Carassius auratus*): Israeli and Kimmel, 1996; Atlantic cod (*Gadus morhua*): Herbert and Steffensen, 2005; rainbow trout: Steffensen and Farrell, 1998; sea bream (*Sparus aurata*): Remen et al., 2015). If locomotor activity represents a complex and situation-specific pattern, any visual increase or decrease of the swimming activity in tanks can be easily observed by fish farmers and thus would represent a relevant indicator of impaired welfare. Stereotypies, jumps or accelerations are usually linked with stressful aquaculture procedures and usually represent a behavioural indicator of poor welfare in farmed fish (Ellis et al., 2002; van de Nieuwegiessen et al., 2008). Startle responses have been observed in cases of severe acute hypoxia when salmonids get to the point of limiting oxygen saturation (LOS, below the threshold for upholding routine metabolism), which is the point where vigorous escape kicks in (Remen et al., 2013). In the present study, we showed that this category of behaviours was under-expressed in the stressed group. The activity level seems to reach a floor effect, probably due to the extent of the stressor, which combined several physic-chemical parameters. Surface activity, including stereotypies and jumps, also decreased in the control group. In this study, the period of acclimation to the experimental conditions lasted for 2 weeks, which is probably not a long enough duration for fish to cope with the new rearing units (size, stocking density). In a previous study, we observed behavioural perturbations 10 days after transferring fish (Colson et al., 2015a). We attributed the global decrease in activity during the acclimation period to the possibly occurring resolution of dominance relationships. The present decreased activity, measured by lower levels of repetitive rubbing and jumping in the controls, confirms the importance of the acclimation period duration in behavioural studies and welfare assessment. This period should last for at least 2 weeks following fish transfer.

The reduced swimming activity observed under degraded water quality was correlated with increased shoaling behaviour. Fish responded to unfavourable conditions by modifying their swimming speed as well as their use of space. Whoriskey *et al* (1985) reported that under hypoxic conditions, three-spined sticklebacks (*Gasterosteus aculeatus*) formed schools as an anti-predator response, rather than attempting to escape. Under hypoxia, salmonids may adjust their spatial positioning within the current velocity gradients in the area with the best water quality, as well in accordance with the influent normoxic water. In the present study, we observed that the latter scenario caused aggregation close to the water inlet. Moreover the water current velocity was decreased in the stressed group, which might also explain the difference in fish position between treatments. Therefore, any modification in fish behaviour (activity and space use) may be easily detectable at group level and represents a first sentinel welfare marker that may be subsequently linked to more subtle sentience-based indicators.

### Open-field test

Chronic stress sometimes results in apathy or blunted emotion, while in other cases it leads to heightened emotional reactivity. Apathy would likely develop when an animal has no way of avoiding negative events, whereas hyper-reactivity would occur when an animal thinks it can control the events (Dantzer and Mormède, 1983; Taghzouti et al., 1999; Boissy et al., 2001; Alcaro et al., 2002; Destrez et al., 2013). Here, we investigated whether either emotional arousal or apathy might be a consequence of a previous chronic stress. However, no significant differences in fear-related behaviour − neither startle responses, nor apathetic behaviour (immobility, thigmotaxis) − appeared between the stressed and the control fish subjected to the open-field test (i.e. social isolation in a novel environment). The flight strategy is frequently observed in chronically stressed animals subjected to an additional acute stressor (mammals: Destrez et al., 2013; birds: Calandreau et al., 2011; zebrafish (*Danio rerio*): Piato et al., 2011) however it was not observed here. One could expect that fish previously exposed to poor water quality might use less energy-demanding forms of response, such as hiding. Complete immobility close to the wall of the tank represents the only available strategy to hide in barren environments for cultured fish. However, immobility and thigmotaxis levels were similar between the stressed and the control fish. Further study is needed to clarify whether the absence of a difference in emotion-related behaviour between the two groups is because the environmental stressor was too mild or because oxygen level is not a major regulator of emotional reactivity. This might also be explained by the standard water quality reversal 24 hours before the test, which could have disrupted the effects. However, this reversal was necessary to avoid any behavioural bias due to neuromuscular inhibition in fish during low oxygen conditions.

Emotional reactivity is inferred from behavioural components and also from neurophysiological processes (cardiac and cortisol reactions). Regarding hypothalamus– pituitary–interrenal (HPI) axis sensitivity, cortisol levels measured 30 minutes after the beginning of the open-field test were higher in fish previously exposed to poor water quality. This is in accordance with previous studies, which also revealed a cortisol increase after a chronic period of hypoxia in Atlantic cod (Herbert and Steffensen, 2005), sturgeon (*Huso huso*) (Lakani et al., 2013) and Arctic char (*Salvelinus alpinus*) (Sandblom et al., 2013). However, in three-spined stickleback, O’Connor et al. (2011) found no measurable effect of a prolonged period (7 days) of hypoxia on cortisol levels, suggesting that these fish acclimated to the low level of oxygen over time. In our study, the chronic stressor lasted for a longer period (3 weeks) and hypoxia was combined with an increase in ammonia nitrogen levels. This suggests that a prolonged period of poor water quality does have long-lasting effects on the HPI axis sensitivity when fish are subjected to an additional acute stress, even after 24 hours of normoxia recovery, and despite the absence of a behavioural response. Similarly, Vindas et al. (2016) observed high cortisol baseline values and strong cortisol responsiveness to acute stress in emaciated and behaviourally inhibited salmon. These authors also reported high baseline serotonergic activity and inability of further response to acute stress in these fish, and proposed this as a behavioural mediator for avoiding further stress exposure.

### Human avoidance test

In practice, individual emotional responses of fish subjected to isolation in a novel tank remain difficult to evaluate in husbandry systems whereas fish group reactivity is more easily detectable. We, therefore, measured fish group behavioural responses to the approach of a human, which can represent a potentially threatening event for farmed fish. During the experiment, fish were automatically fed and we therefore did not expect fish to have a positive association with a human presence (Speare et al., 1995). As expected, in the control group we observed strong human avoidance, with fish grouping as a shoal at the opposite side of the tank, when the experimenter approached. However, human avoidance was significantly lower in stressed fish than in controls. This result suggests that chronic stress leads to an inhibition of the normal escaping behaviour classically expressed in automatically fed fish. This inhibited human avoidance in the chronically stressed fish possibly reflects global apathy, a common expression of poor welfare as described above (Boissy et al., 2001). This might also be linked to the decrease in swimming activity previously observed in various fish species exposed to hypoxia conditions (Steffensen and Farrell, 1998; Remen et al., 2015).

This finding validates the relevance of our human avoidance test, an adaptation of a classical test commonly used in terrestrial farmed animals (see Waiblinger et al. (2006) for a review). To our knowledge, this is the first time that reactivity to human presence has been reported in farmed fish. The major advantage of the human avoidance test is that it can be easily applied on farms at the group level and can be considered to be a sentience-based indicator of welfare.

### Appetitive conditioning

Fish from the control group, but not the stressed group, correctly learned to associate a neutral stimulus with a food reward. In the control group, the conditioned response tended to be higher than baseline level after 8 trials and it was higher in the control than in the stressed group at trial 5, showing the remarkable learning skills of the rainbow trout in a classical appetitive conditioning paradigm. The acquisition is faster than previously observed in sea bream (*Sparus aurata*), which associate a US with a CS after approximately 16 trials (Folkedal et al., 2018). The inherent stress effect of a reduced appetite, as demonstrated by a significant reduction in growth in the stressed group compared to the control group, might represent an explanation of the impaired learning in the stressed group. Nevertheless, fish from both groups did finally group around the feeder when the food was effectively distributed at the end of trial 18. This final grouping excludes a putative decrease in physical capacities induced by hypoxia, since stressed fish were still able to swim and orientate towards the food when the distribution did occur. The lack of responsiveness when the CS was activated alone suggests a failure to adapt to the behavioural challenge proposed by the rewarded conditioning. With regards to adaptive function, fish previously exposed to poor water quality have to allocate their energy to vital functions avoiding useless displacements, here represented by a non-immediately rewarded signal. Interestingly, this effect was observed even 2 weeks after the end of the hypoxia and ammonia exposure, which suggests that such a chronic stressor has profound and long-lasting effects on food-anticipatory behaviour. More than a learning skill assessment, the inclination of an individual to expect a positive event is by itself a relevant way to measure a positive emotional state (Spruijt et al., 2001). Here, the lack of this inclination might contrarily indicate a global apathy in stressed fish. An inhibition of appetitive conditioned responses was also demonstrated in Atlantic salmon (*Salmo salar*) a few hours after different acute stressors (Folkedal et al., 2012b), but this is the first time such an inhibition has been demonstrated 2 weeks after a chronic exposure to poor water quality in rainbow trout. In another study including different feed rations, (Folkedal et al., 2012a) demonstrated that hunger level was a mediator between the conflicting motivational systems in salmon post-smolt. The conditioned response towards food appears to be a trade-off between competitive motivational systems (hunger, fear, etc.) where the decision making of stressed fish is directed towards stress coping as well as influenced by a general lack of feeding motivation. Here, there is still a discrepancy between the appetitive response and feed intake, since the fish may feed but not respond strongly to the CS. The highly responsive HPI axis observed in chronically stressed fish might have long-term effects on brain function, possibly due to the dysfunction of stress-related neurotransmitters (serotonin) that can interfere with efficient storage of information (Mendl et al., 2001; Lepage et al., 2005; Laursen et al., 2013; Vindas et al., 2016). Further studies would be required to measure neurotransmitter levels when rainbow trout are exposed to poor water quality and to confirm their possible implication in appetitive conditioning, even 2 weeks after the end of hypoxia and ammonia exposure. To rule out appetite as a key factor of poor water quality challenge or not, an alternative paradigm would be to use an aversive US, as already tested in rainbow trout (Yue et al., 2004). However, fish food-anticipatory behaviour is an easy and applicable way to measure a possible negative effect of chronic stress in aquaculture systems. Farmers can easily observe food-anticipatory behaviour during routine inspection before food distribution and the present study demonstrates that it provides a good operational sentience-based welfare indicator at the group level. Now commonly used in terrestrial farmed animals (see Paul et al. (2005) for a review), cognitive science also seems to offer novel and promising approaches for welfare assessment in fish.

## 5. Conclusion

Accurate assessment of emotion and cognition is now crucial in animal welfare research. This study provides fresh insights, which can be easily adapted for farmed fish. Here we show that any behavioural modifications observed at the group level could be considered to be sentinel or “Iceberg” indicators providing early warning in case of environmental disturbance. Then practical sentience-based tests can be performed in fish kept in their home tank. We demonstrated that behavioural responses to a standardized human approach and food distribution anticipation could be directly observed by farmers and considered as sensitive welfare markers. These innovative sentience-based tools should be of great interest to the European aquaculture industry, which should be able to control fish welfare by taking into account the mental states of the animals.

## Acknowledgements

We thank J Bobe for his careful reading of the manuscript and C Cheung for her English advices. This work received funding from the European Union Seventh Framework Program (FP7/2007–2013) under grant agreement No. 262336, AQUAEXCEL.

